# A hybrid approach for predicting transcription factors

**DOI:** 10.1101/2022.07.13.499865

**Authors:** Sumeet Patiyal, Palak Tiwari, Mohit Ghai, Aman Dhapola, Anjali Dhall, Gajendra P. S. Raghava

## Abstract

Transcription factors (TFs) are essential DNA-binding proteins that regulate the rate of transcription of several genes and controls the expression of genes inside a cell. The prediction of TFs with high precision is important for understanding number of biological processes such as cell-differentiation, intracellular signaling, cell-cycle control. In this study, we developed a hybrid method that combine alignment-based and alignment-free methods for predicting transcription factors with higher accuracy. All models have been trained, tested and evaluated on a large dataset that contain 19406 TFs and 523560 non-TFs protein sequences. In order to avoid biasness in evaluation, dataset is divided in training and validation/independent dataset, where 80% data was used for training and remaining 20% for external validation. In case of alignment-free methods, models are developed based on machine learning techniques using compositional features of a protein. Our best alignment-free model obtained AUC 0.97 on independent dataset. In case of alignment-based method, we used BLAST at different cut-off to predict transcription factors. Though alignment-based method shows excellent performance but unable to cover all transcription factor due to no-hits. In order to combine power of both, we developed a hybrid method that combine alignment-free and alignment-based method; achieved maximum AUC of 0.99 on independent dataset. The method proposed in this study perform better than existing methods. We incorporated the best models in the webserver/standalone package “TransFacPred” (https://webs.iiitd.edu.in/raghava/transfacpred).

**Key Points:**

- Transcription factors (TFs) are vital DNA-binding proteins.
- A hybrid method for the prediction of TFs using sequence information.
- Computer-aided model were developed using machine-learning algorithm to predict TFs.
- Alignment-based and alignment-free approaches were used for the prediction.
- A user-friendly webserver, python- and Perl-based standalone package available.

## Introduction

Transcription factors (TF) are DNA-binding proteins that bind to specific DNA segments to control the expression of the genes [1-3]. These TFs or regulators control specific cell types, cell differentiation, gene regulatory pathways, and immune responses [4-6]. Recognition of TFs is the first step for understanding the transcription regulatory system [7]. Mis-regulation and mutations in TFs or their binding regions lead to the development of several disorders like Rubinstein-Taybi, CHOPS syndromes, Coffin-Siris, etc. [4, 8-10]. Moreover, several biological mechanisms such as chromosomal translocation, aberrant gene expression, point substitutions, and mutations associated with the non-coding DNA result in the alteration of transcription factor binding sites in various cancer types [11-15]. In addition, several inflammatory, autoimmune diseases and improper immune development are associated with the mis-regulation of NF-kB transcription factor [16]. Studies also reveal that, with a better understanding of the transcriptional regulations, one can control gene expression in various genetic perturbations [4, 17, 18]. In the past, several attempts in clinical have been made to target/inhibit or modulate transcription factor DNA-binding activity in disease conditions [19-21].

In addition, with the availability of enormous genome sequencing data, many methods have been developed to identify TFs [22]. It is not practically feasible to identify TFs in genomics using experimental techniques. In order to overcome these limitation, number of in silico methods have been developed to annotate TFs at genome scale [23]. Zheng et al. developed an approach to predict different classes of TFs like Helix-turn-helix, Beta-scaffold and zinc-coordinating DNA binding domains [24]. Eichner et al. developed a four-step workflow for identifying a DNA-binding domains and inferring the DNA motif in a protein. TFPredict uses machine/deep learning techniques to predict a transcription factor [25]. Another tool BART has been developed to predict functional factors that bind at cis-regulatory regions from gene list or a ChIP-seq dataset [26]. Recently, Kim et al. constructed a deep neural network method named DeepTFactor to predict predicting real-valued TF binding intensities, whether TF-DNA binding or unbinding [7]. In this study, they created and used largest possible dataset for developing an accurate and reliable method. The available methods are computationally expensive and require the domain knowledge.

In order to overcome limitations of existing methods, we developed a improved method for predicting transcription factor with high accuracy. Initially, we developed homology or alignment-based methods for prediction. These alignment-based methods exhibit high performance if query TF have high similarity with target TF in database. These methods fail if a query TF have poor similarity with known TFs in database or high similarity with non-TF. In order to overcome these limitations, we developed alignment-based method. In case of alignment-based methods, different machine learning techniques have been used for building prediction models. In these methods, composition of TFs is used as input feature for building prediction models. In order to combine power of both alignment free and alignment-based methods, we developed a hybrid method. In order to facilitate scientific community, we developed a webserver and standalone software package TransFacPred which is available https://webs.iiitd.edu.in/raghava/transfacpred and https://github.com/raghavagps/transfacpred

## Materials and Methods

### Dataset Collection and pre-processing

We obtained the TF and non-TF protein sequence dataset from UniProt Knowledge-base (UniProtKB)/Swiss-Prot database [27, 28], which was released in September 2019. The dataset was parsed and classified into TF and non-TFs using the Gene Ontology (GO) annotation. The complete table for GO terms used to classify the TFs and non-TFs is shown in Supplementary Table S1. Here, we obtained a total of 21802 TF sequences and 539374 non-TF sequences. We removed the redundant sequences and sequences with the non-natural amino acids from the TF and non-TF datasets. For the positive dataset we obtained 19406 unique TF sequences out of 21802 sequences. In case of negative dataset, we obtained 523560 non-TF sequences from 539374 entries. The final dataset comprises 19406 TFs (positive) and 523560 non-TFs (negative) protein sequences. Then, we followed the standards used in previous studies [29, 30] and split the whole dataset into 80% training dataset comprising of 434373 sequences (15525 TFs and 418848 non-TFs), and 20% validation dataset containing 108594 sequences (3882 TFs and 104712 non-TFs).

### Feature Generation

#### Composition-based features

Pfeature [31] used in this study to computed amino acid composition (AAC) and di-peptide composition (DPC) based features of positive and negative datasets. In case of AAC, a feature vector of length 20 was generated which represents the composition of 20 amino acids in the sequence. Dipeptide composition is used to encapsulate the global information about each sequence, which gives a fixed vector of length 400 (20 × 20).

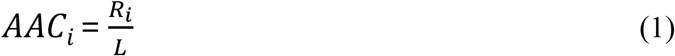

where ***AAC***_***i***_ is amino acid composition of residue type ***i***; ***R***_***i***_ and ***L*** number of residues of type ***i*** and length of sequence, respectively.

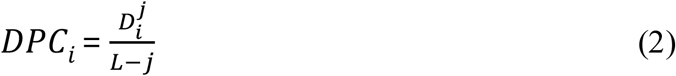

Where ***DPC***^***j***^_***i***_ is the fraction or composition of dipeptide of type ***i*** for jth order. ***D***^***j***^_***i***_ and ***L*** are the number of dipeptides of type ***i*** and length of a protein sequence, respectively.

### One-hot encoding (OHE)

We have implemented one hot encoding approach for feature generation using TF and non-TF sequences. It is a representation of categorical variables as binary vectors. First it requires the categorical values be mapped to integer values. Then, each integer value is represented as a binary vector that is all zero values except the index of the integer, which is marked with a 1. In OHE, each amino acid is represented by the vector size of length 21, for instance, A is described as 1,0,0,0,0,0,0,0,0,0,0,0,0,0,0,0,0,0,0,0,0; which consist of 20 natural amino acids and one dummy variable, whereas X is represents as 0,0,0,0,0,0,0,0,0,0,0,0,0,0,0,0,0,0,0,0,0.

### Model development and Cross-Validation

In this study, we have implemented a number of classifiers in order to develop prediction models. Here, we have used Scikit-learn based traditional machine learning algorithms such as Decision Tree (DT), eXtreme Gradient Boosting (XGB), Random Forest (RF), Gaussian Naive Bayes (GNB), K-Nearest Neighbor (KNN), Extra Tree (ET), Logistic Regression (LR), and Support Vector Classifier (SVC). To avoid the curse of biasness and overfitting of models, we have performed five-fold cross-validation on the training dataset [32-34]. In this approach, the training dataset is stratified into five sets, where the model is trained on four sets and tested on the remaining one. The same process is repeated five times in such a way that each set get to be act as testing dataset. The final performance is the average of performances resulted from each iteration.

### Similarity Search approach

In this study, we have also implemented similarity search using BLAST [35] as it is a widely used tool to annotate the sequences. We have used it to classify the sequences as transcription factor or non-transcription factor based on their similarity. The blastp suite of NCBI-BLAST+ version 2.2.29 was used to perform the similarity search. Training dataset was used to create the custom database, makeblastdb suite of NCBI-BLAST+ was used for the same. Sequences in the validation dataset were hit against the custom database to assign the class as transcription factor or non-transcription factor based on their similarity with the sequences in the database. In the present study, we have considered the top-hit of BLAST to assign the classes, such that if the top-hit of the BLAST is against transcription factor sequence of the database, then the protein is assigned as transcription factor otherwise non-transcription factor. We have run the BLAST at different e-value cut-offs varying from 1e-6 to 1e+3, in order to find the optimal value to classify the transcription factors.

### Performance Evaluation

We have used various performance evaluation parameters such as accuracy, sensitivity, specificity, F1-score, Area Under Receiver Operating Characteristics (AUC), and Matthews Correlation Coefficient (MCC). Where, sensitivity (see equation 1), specificity (see equation 2), accuracy (see equation 3), F1-score (see equation 4), and MCC (see equation 5) are threshold-dependent parameters. On the other side, AUC is threshold-independent parameters. The equations of various performance evaluation parameters are provided below.

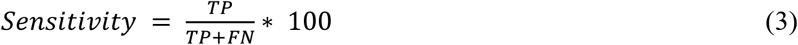

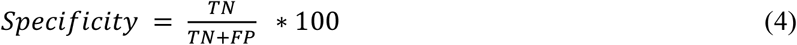

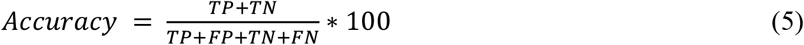

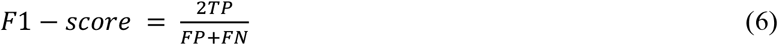

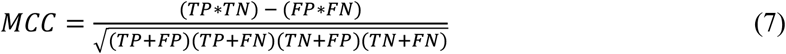

Where, FP is false positive, FN is false negative, TP is true positive and TN is true negative.

## Results

### Compositional Analysis

In this study, we have performed the amino acid based compositional analysis for TF, non-TF, and general proteome, in order to compare the abundance of the residues in these classes. Figure 1 represents the average percent composition of each residue in proteins belongs to TF and non-TF class, and compared the same with average percent composition of general proteome derived from Swiss-Prot database. As exhibit by the bar-plot, transcription factors are rich in E, P, Q, R and S residues as compared to the non-transcription factors, whereas residue A, G, I, and V are abundant in non-transcription factor proteins.

**Figure 1:**
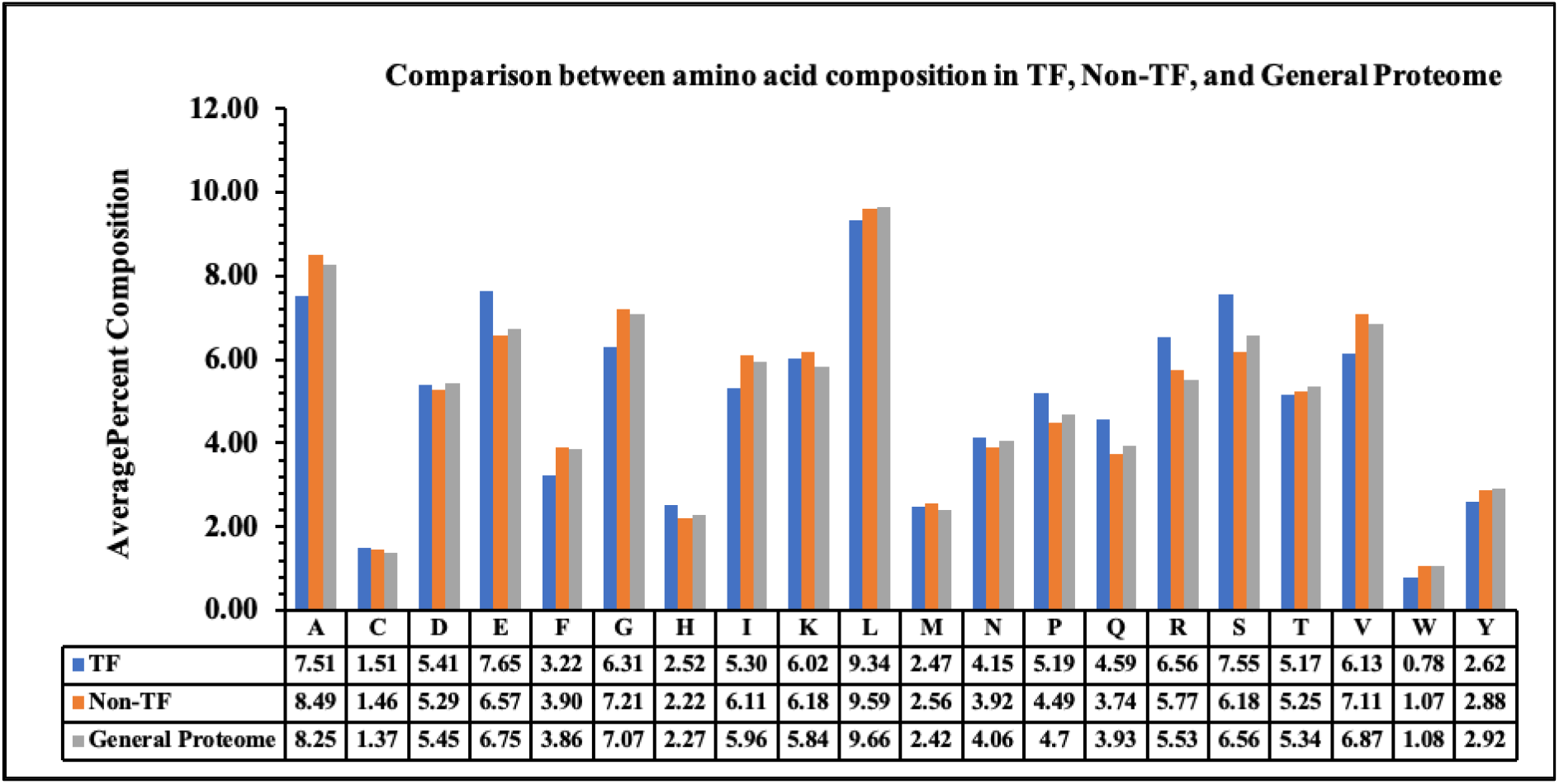
Average percent composition of amino acid residues in TF, Non-TF, and General Proteome.

### Performance on alignment-based method

In order to classify the transcription factors using alignment-based method, we have performed the similarity search using BLAST by varying e-value from 1.00E-06 to 1.00E+03. In this approach, we have created the database using the sequences in the training dataset, and hit the query proteins in the validation dataset against it, and considered the top-hit to assign the class to each query proteins. The performance at each value is reported in Table 1. As shown in Table 1, BLAST has achieved a good performance for predicting the transcription factors but cannot cover the entire dataset. Moreover, as e-value is increasing, the probability of correct prediction is decreasing. Hence, BLAST alone is not sufficient for predicting the transcription factors.

**Table 1:** Performance on alignment-based approach at different e-values.

| E-Value | No hits [Positive] | Probability of Correct Prediction |
| --- | --- | --- |
| 1.00E-06 | 68 | 95.44 |
| 1.00E-05 | 57 | 95.40 |
| 1.00E-04 | 44 | 95.28 |
| 1.00E-03 | 39 | 95.29 |
| 1.00E-02 | 32 | 95.30 |
| 1.00E-01 | 29 | 95.28 |
| 1.00E+00 | 22 | 95.26 |

### Performance on alignment-free method

We have implemented eight traditional machine learning classifiers such as, DT, RF, LR, XGB, GNB, KNN, ET, and SVC using various features like AAC, DPC, and AAC+DPC as the input feature, to classify the protein sequences into TFs and non-TFs. We have trained out the model on the 80% training dataset and evaluate the performance on the 20% remaining dataset. First, we have developed various prediction models using amino acid composition, and the performance of each classifier is reported in Table 2. As shown by Table 2, ET-based model outperforms the other models with AUC 0.97 on training and independent dataset, with balanced sensitivity and specificity.

**Table 2:** Performance of various classifiers using AAC as input feature.

| Classifier | Training Dataset |  |  |  |  |  |  | Independent Dataset |  |  |  |  |  |  |
| --- | --- | --- | --- | --- | --- | --- | --- | --- | --- | --- | --- | --- | --- | --- |
|  | Sens | Spec | Acc | AUC | F1 | K | MCC | Sens | Spec | Acc | AUC | F1 | K | MCC |
| **DT** | 52.435 | 98.192 | 96.557 | 0.753 | 0.523 | 0.505 | 0.505 | 52.486 | 98.273 | 96.637 | 0.754 | 0.529 | 0.511 | 0.511 |
| **RF** | 91.028 | 89.057 | 89.127 | 0.964 | 0.708 | 0.698 | 0.701 | 91.961 | 89.195 | 89.294 | 0.968 | 0.721 | 0.711 | 0.713 |
| **LR** | 73.895 | 74.756 | 74.725 | 0.814 | 0.214 | 0.168 | 0.215 | 74.130 | 75.001 | 74.970 | 0.813 | 0.213 | 0.169 | 0.215 |
| **XGB** | 85.592 | 86.791 | 86.748 | 0.940 | 0.582 | 0.568 | 0.574 | 86.988 | 86.966 | 86.967 | 0.946 | 0.589 | 0.575 | 0.583 |
| **KNN** | 84.684 | 94.997 | 94.628 | 0.913 | 0.671 | 0.661 | 0.669 | 85.803 | 95.011 | 94.682 | 0.919 | 0.674 | 0.663 | 0.672 |
| **GNB** | 67.835 | 71.707 | 71.569 | 0.772 | 0.235 | 0.202 | 0.206 | 67.070 | 72.002 | 71.825 | 0.767 | 0.235 | 0.201 | 0.206 |
| **ET** | 90.461 | 90.866 | 90.852 | 0.967 | 0.733 | 0.724 | 0.729 | 91.033 | 90.949 | 90.952 | 0.968 | 0.745 | 0.736 | 0.740 |
| **SVC** | 80.246 | 80.812 | 80.791 | 0.891 | 0.488 | 0.469 | 0.470 | 81.474 | 81.186 | 81.196 | 0.897 | 0.489 | 0.469 | 0.470 |

Similarly, various machine learning models were developed to classify TFs using DPC as the input features. Table 3 represents the performance of models based on each classifier, and model based on XGB classifier performed best among the other classifiers with AUC of 0.96 on training and validation dataset.

**Table 3:** Performance of various classifiers using DPC as input feature.

| Classifier | Training Dataset |  |  |  |  |  |  | Independent Dataset |  |  |  |  |  |  |
| --- | --- | --- | --- | --- | --- | --- | --- | --- | --- | --- | --- | --- | --- | --- |
|  | Sens | Spec | Acc | AUC | F1 | K | MCC | Sens | Spec | Acc | AUC | F1 | K | MCC |
| **DT** | 52.544 | 98.180 | 96.549 | 0.754 | 0.522 | 0.505 | 0.505 | 52.409 | 98.193 | 96.557 | 0.753 | 0.522 | 0.504 | 0.504 |
| **RF** | 90.648 | 89.220 | 89.271 | 0.964 | 0.720 | 0.710 | 0.715 | 90.518 | 89.444 | 89.482 | 0.964 | 0.728 | 0.719 | 0.726 |
| **LR** | 80.343 | 80.711 | 80.698 | 0.876 | 0.301 | 0.265 | 0.303 | 80.752 | 80.779 | 80.778 | 0.878 | 0.308 | 0.272 | 0.309 |
| **XGB** | 90.113 | 90.305 | 90.298 | 0.965 | 0.720 | 0.710 | 0.715 | 90.054 | 90.505 | 90.488 | 0.966 | 0.720 | 0.711 | 0.716 |
| **KNN** | 84.117 | 96.321 | 95.885 | 0.913 | 0.714 | 0.703 | 0.704 | 83.767 | 96.225 | 95.780 | 0.912 | 0.719 | 0.709 | 0.709 |
| **GNB** | 75.866 | 49.497 | 50.439 | 0.694 | 0.166 | 0.122 | 0.135 | 76.269 | 49.668 | 50.619 | 0.696 | 0.165 | 0.122 | 0.133 |
| **ET** | 90.938 | 88.708 | 88.788 | 0.965 | 0.757 | 0.749 | 0.752 | 90.673 | 88.889 | 88.952 | 0.964 | 0.756 | 0.747 | 0.753 |
| **SVC** | 88.864 | 92.169 | 92.051 | 0.960 | 0.781 | 0.774 | 0.778 | 89.307 | 92.171 | 92.069 | 0.964 | 0.787 | 0.779 | 0.782 |

In the next step, we have combined the AAC and DPC feature which resulted into the vector of size 420 for each protein and developed prediction models. We have used eight different classifiers and their performance is reported in Table 4. Similar to the performance on individual features, XGB-based model has performed the best among all the other classifiers with AUC of 0.97 on training and independent dataset.

**Table 4:**
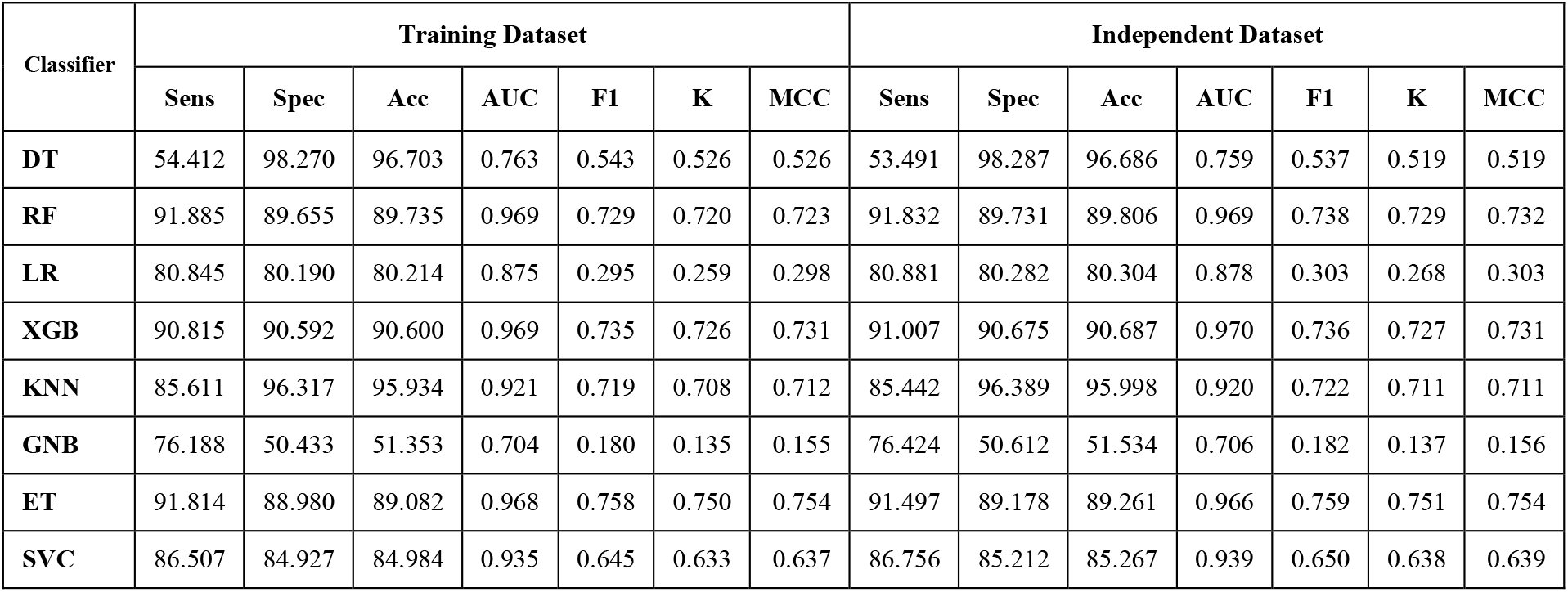
Performance of various classifiers using combination of AAC and DPC as input feature.

### Performance of deep learning models

We have also developed the deep learning technique-based prediction models to classify the TFs using different features such as AAC, DPC, AAC+DPC, and one-hot encoding (OHE). Table 5 exhibits the performance of different models on validation dataset using different features. As shown by the Table 5, CNN based model with one-hot encoding as the input feature performed best with AUC 0.95 on the independent dataset.

**Table 5:** Performance of deep learning model (CNN) using various features on independent dataset.

| Feature | Sensitivity | Specificity | Accuracy | AUROC | F1 | MCC |
| --- | --- | --- | --- | --- | --- | --- |
| AAC | 8.00 | 99.00 | 96.30 | 0.54 | 0.14 | 0.24 |
| DPC | 53.22 | 99.73 | 97.92 | 0.76 | 0.67 | 0.68 |
| AAC+DPC | 59.34 | 99.49 | 97.93 | 0.79 | 0.69 | 0.69 |
| OHE | 91.27 | 98.61 | 98.32 | 0.95 | 0.81 | 0.81 |

### Performance of hybrid (alignment-based + alignment-free) model

We have also developed the hybrid to classify the TFs in which we used the similarity search using BLAST to make the prediction. In hybrid approach, we have combined the outputs from ET-based model developed using amino acid composition and BLAST search, to make the final prediction. Table 6 exhibits the performance of hybrid model at different e-values on validation dataset. As shown by the Table 6, at each e-value the AUC achieved the AUC 0.99, with balanced sensitivity and specificity, in terms of accuracy e-value 1.00E+02 attained the maximum value of 97.013%. This model has been incorporated in the backend of the server TransFacPred to predict if the submitted protein is a TF or non-TF.

**Table 6:** Performance of hybrid method (AAC + BLAST) on independent dataset.

| **E-value** | **Sensitivity** | **Specificity** | **Accuracy** | **AUC** | **F1** | **K** | **MCC** |
| --- | --- | --- | --- | --- | --- | --- | --- |
| **1.00E-06** | 95.88 | 95.41 | 95.42 | 0.99 | 0.94 | 0.93 | 0.93 |
| **1.00E-05** | 95.83 | 95.56 | 95.57 | 0.99 | 0.94 | 0.93 | 0.93 |
| **1.00E-04** | 95.70 | 95.76 | 95.76 | 0.99 | 0.94 | 0.94 | 0.94 |
| **1.00E-03** | 95.70 | 95.93 | 95.92 | 0.99 | 0.94 | 0.94 | 0.94 |
| **1.00E-02** | 96.03 | 96.03 | 96.03 | 0.99 | 0.94 | 0.94 | 0.94 |
| **1.00E-01** | 96.16 | 96.18 | 96.18 | 0.99 | 0.94 | 0.94 | 0.94 |
| **1.00E+00** | 96.34 | 96.41 | 96.41 | 0.99 | 0.94 | 0.94 | 0.94 |
| **1.00E+01** | 96.96 | 96.76 | 96.77 | 0.99 | 0.93 | 0.93 | 0.93 |
| **1.00E+02** | 97.06 | 97.01 | 97.01 | 0.99 | 0.93 | 0.92 | 0.92 |
| **2.00E+02** | 97.06 | 97.15 | 97.15 | 0.99 | 0.93 | 0.92 | 0.92 |
| **1.00E+03** | 97.06 | 97.24 | 97.24 | 0.99 | 0.93 | 0.92 | 0.92 |

### Comparison with DeepTFactor

In order to understand the advantages or disadvantages of the new methods, it is crucial to compare the same with the existing methods. Hence, we have compared the performance of our models with the recently published method DeepTFactor [7]. We have evaluated our and DeepTFactor models on the validation dataset and as signified in Table 7, our model has performed better in terms of each parameter.

**Table 7:**
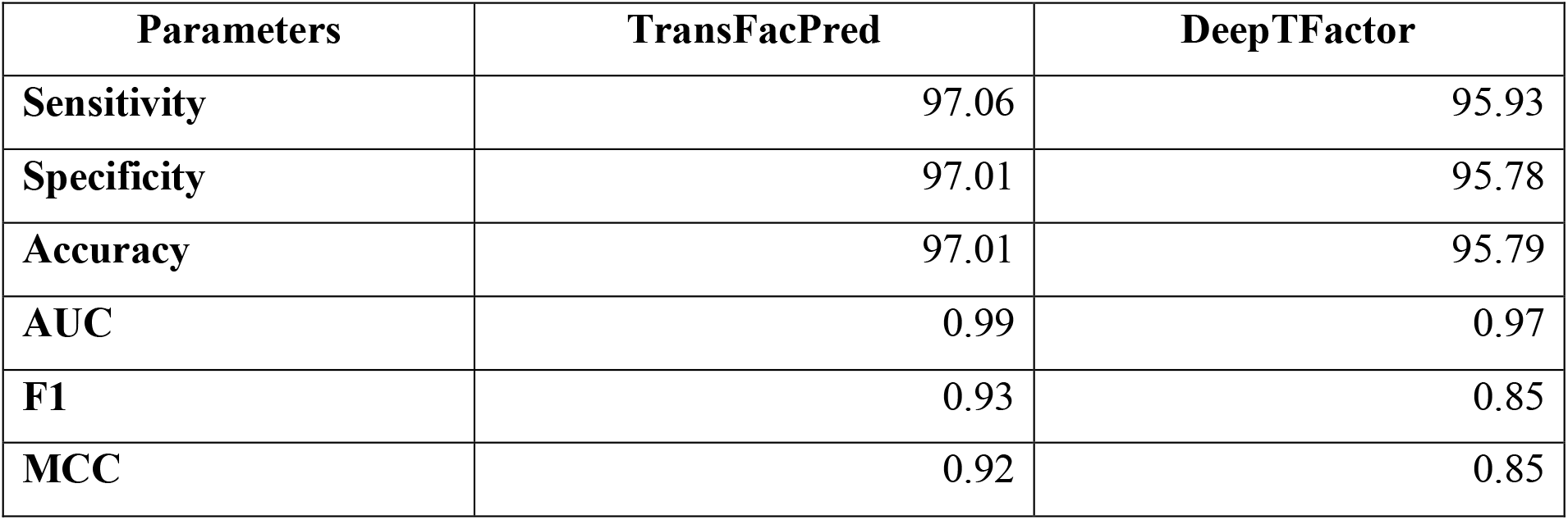
Comparison of performance of our best model with DeepTFactor on independent dataset.

In addition, we have also compared the processing time of DeepTFactor and TransFacPred standalone using ML and hybrid model, by providing various number of sequences at a time and found that as the number of sequences increases DeepTFactor takes longer time as shown in Table 8. On the other hand, we have implemented AAC based machine learning model as well as hybrid model and compared the performance. ML based model took less time as compare to DeepTFactor with equivalent AUC, whereas hybrid model performed best but took more time to provided output.

**Table 8:** Comparison between the processing time of DeepTFactor and TransFacPred.

| Number of Sequences | Method | Time (in seconds) |  |  |
| --- | --- | --- | --- | --- |
|  |  | Real | User | System |
| 50 | DeepTFactor | 13.285 | 3.882 | 1.188 |
|  | TransFacPred [ML] | 7.666 | 1.551 | 0.998 |
|  | TransFacPred [Hybrid] | 24.111 | 22.079 | 1.254 |
| 1000 | DeepTFactor | 55.201 | 51.37 | 3.954 |
|  | TransFacPred [ML] | 37.208 | 2.649 | 1.157 |
|  | TransFacPred [Hybrid] | 436.071 | 429.062 | 3.157 |
| 108594 | DeepTFactor | 6014.113 | 5629.047 | 375.138 |
|  | TransFacPred [ML] | 134.387 | 130.191 | 1.945 |
|  | TransFacPred [Hybrid] | 47932.78 | 47583.942 | 304.83 |

### Web-Server Implementation

We have developed easy to use webserver TransFacPred and a standalone package. Our web server has two major modules; Predict and BLAST Search. Predict module allows the users to predict TFs using alignment-free method or hybrid method (see Figure 2). BLAST search module allows users to perform blast search against database of TFs and non-TFs used in this study. In addition to webserver, we have also developed standalone package in Python. This package is suitable for scanning TFs at genome scale where user can run this package on their local machine.

**Figure 2:**
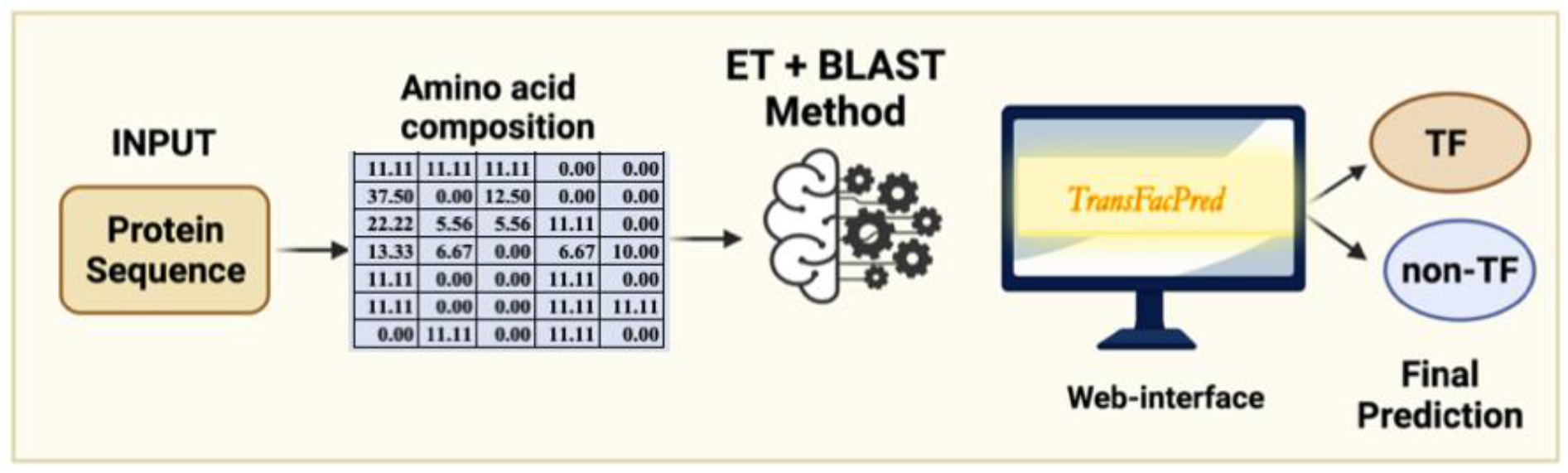
Graphical representation for the working of TransFacPred web-server using hybrid model.

## Discussion

TFs initiate the process of transcription and hence plays a major role in deciding the fate of a cell or cellular process [36, 37]. Identification of novel or unknown TFs using experimental based techniques such as RNA sequencing (RNA-seq) and Chromatin Immunoprecipitation sequencing (ChIP-seq) experiment is a tiring and cost expensive task [38]. Previously, a number of methods have been developed for the prediction of TFs [7, 24, 25]. In order to facilitate the researchers working in this field, we have made a systematic attempt to develop a highly accurate method with the capability of classifying TFs using the primary sequence information. Based on GO terms, sequences from UniProtKB/Swiss-Prot were assigned as either TF or non-TF. At first, there was total of 561176 sequences, out of which 21802 were TFs and 539374 were non-TFs, after preprocessing the datasets the final dataset was comprised of 19406 TFs and 523560 non-TFs.

The preliminary composition analysis on this dataset showed that the TFs are rich in E, P, Q, R, and S amino acids. Further, sequence-based features were computed using Pfeature software and various machine learning techniques were implemented to exploit their capabilities to classify the sequences as either TFs or non-TFs. Our models were trained on 80% of the dataset using different set of features and validated on the remaining previously unseen 20% dataset. We obtained an AUC of 0.96 on training as well as validation dataset using amino acid composition-based features. Out of all the models, hybrid model which is the combination of ET-based model developed on amino acid composition and BLAST search performed best with AUC 0.99 on the validation dataset with balanced sensitivity and specificity. We have also compared our method with the recently published method DeepTFactor. We have trained our models on the training dataset and evaluate the performance of DeepTFactor model and our models on the validation dataset, and found out that models of TransFacPred has outperformed DeepTFactor in terms of AUC and other parameters We anticipate that this research will aid the researchers working in the field of genomics and proteomics. Figure 3 represents the complete flow of this study.

**Figure 3:**
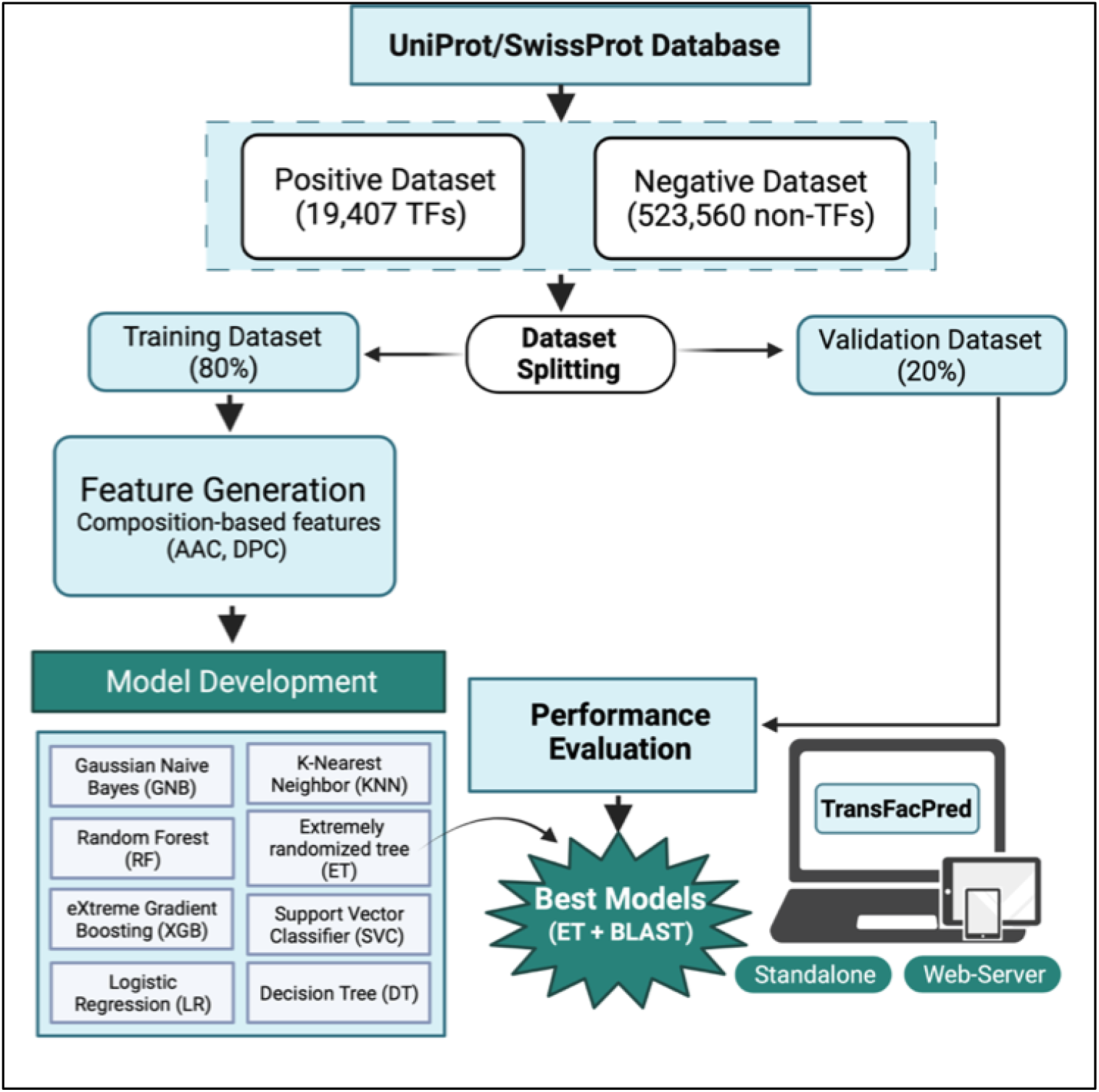
Complete workflow for TransFacPred.

## Acknowledgements

Authors are thankful to the Department of Biotechnology (DBT) and Department of Science and Technology (DST-INSPIRE) for fellowships and the financial support. Authors are also thankful to Department of Computational Biology, IIITD New Delhi for infrastructure and facilities.

## Authors’ contributions

SP, PT, MG, ADO, and GPSR collected and processed the datasets. SP, PT, MG, and GPSR implemented the algorithms and developed the prediction models. SP, PT, MG, ADO, AD, and GPSR analyzed the results. SP created the back-end of the web server and AD created the front-end user interface. SP, AD, and GPSR penned the manuscript. GPSR conceived and coordinated the project. All authors have read and approved the final manuscript.

## Author’s Biography

1. Sumeet Patiyal is currently working as Ph.D. in Computational Biology from Department of Computational Biology, Indraprastha Institute of Information Technology, New Delhi, India.
2. Palak Tiwari is pursuing M. Tech. in Computer Science and Engineering from Department of Computer Science and Engineering, Indraprastha Institute of Information Technology, New Delhi, India.
3. Mohit Ghai is pursuing M. Tech. in Computer Science and Engineering from Department of Computer Science and Engineering, Indraprastha Institute of Information Technology, New Delhi, India.
4. Aman Dhapola is pursuing M. Tech. in Computer Science and Engineering from Department of Computer Science and Engineering, Indraprastha Institute of Information Technology, New Delhi, India.
5. Anjali Dhall is currently working as Ph.D. in Computational Biology from Department of Computational Biology, Indraprastha Institute of Information Technology, New Delhi, India.
6. Gajendra P. S. Raghava is currently working as Professor and Head of Department of Computational Biology, Indraprastha Institute of Information Technology, New Delhi, India.

