## Supplementary Table S1 for "A hybrid approach for predicting transcription factors"

**Mailing Address of Authors**

Sumeet Patiyal:

Anjali Dhall:

Palak Tiwari:

Mohit Ghai:

Aman Dhapola:

Gajendra P. S. Raghava:

***Corresponding Author**

Prof. Gajendra P. S. Raghava

Head and Professor

Department of Computational Biology

Indraprastha Institute of Information Technology, Delhi

Okhla Industrial Estate, Phase III, (Near Govind Puri Metro Station)

New Delhi, India – 110020

Office: A-302 (R&D Block)

Website: <http://webs.iiitd.edu.in/raghava/>

**Supplementary Figure S1: GO terms used to classify TF sequences and non-TF sequences**

| **Type** | **GO Terms** | **Description** |
| --- | --- | --- |
| Transcription Factor | GO:0000976 | transcription regulatory region sequence-specific DNA binding |
| Transcription Factor | GO:0000977 | RNA polymerase II transcription regulatory region sequence-specific DNA binding |
| Transcription Factor | GO:0000978 | RNA polymerase II cis-regulatory region sequence-specific DNA binding |
| Transcription Factor | GO:0000979 | RNA polymerase II core promoter sequence-specific DNA binding |
| Transcription Factor | GO:0000981 | DNA-binding transcription factor activity, RNA polymerase II-specific |
| Transcription Factor | GO:0000984 | bacterial-type RNA polymerase transcription regulatory region sequence-specific DNA binding |
| Transcription Factor | GO:0000985 | bacterial-type RNA polymerase core promoter sequence-specific DNA binding |
| Transcription Factor | GO:0000986 | bacterial-type cis-regulatory region sequence-specific DNA binding |
| Transcription Factor | GO:0000987 | cis-regulatory region sequence-specific DNA binding |
| Transcription Factor | GO:0000992 | RNA polymerase III cis-regulatory region sequence-specific DNA binding |
| Transcription Factor | GO:0000995 | RNA polymerase III general transcription initiation factor activity |
| Transcription Factor | GO:0001046 | core promoter sequence-specific DNA binding |
| Transcription Factor | GO:0001163 | RNA polymerase I transcription regulatory region sequence-specific DNA binding |
| Transcription Factor | GO:0001164 | RNA polymerase I core promoter sequence-specific DNA binding |
| Transcription Factor | GO:0001165 | RNA polymerase I cis-regulatory region sequence-specific DNA binding |
| Transcription Factor | GO:0001216 | DNA-binding transcription activator activity |
| Transcription Factor | GO:0001227 | DNA-binding transcription repressor activity, RNA polymerase II-specific |
| Transcription Factor | GO:0003700 | DNA-binding transcription factor activity |
| Transcription Factor | GO:0034246 | mitochondrial sequence-specific DNA-binding transcription factor activity |
| Transcription Factor | GO:0098531 | ligand-activated transcription factor activity |
| Transcription Factor | GO:0106250 | DNA-binding transcription repressor activity, RNA polymerase III-specific |
| Transcription Regulation | GO:0001228 | DNA-binding transcription activator activity, RNA polymerase II-specific |
| Transcription Regulation | GO:0006351 | transcription, DNA-templated |
| Transcription Regulation | GO:0006355 | regulation of transcription, DNA-templated |
| Transcription Regulation | GO:0043433 | negative regulation of DNA-binding transcription factor activity |
| Transcription Regulation | GO:0045892 | negative regulation of transcription, DNA-templated |
| Transcription Regulation | GO:0045893 | positive regulation of transcription, DNA-templated |
| Transcription Regulation | GO:0051090 | regulation of DNA-binding transcription factor activity |
| Transcription Regulation | GO:0051091 | positive regulation of DNA-binding transcription factor activity |
| Transcription Regulation | GO:2000142 | regulation of DNA-templated transcription, initiation |
| Transcription Regulation | GO:2000143 | negative regulation of DNA-templated transcription, initiation |
| Transcription Regulation | GO:2000144 | positive regulation of DNA-templated transcription, initiation |
| DNA Binding | GO:0003677 | DNA binding |
| DNA Binding | GO:0008301 | DNA binding, bending |
| DNA Binding | GO:0043565 | sequence-specific DNA binding |
| DNA Binding | GO:0050692 | DNA binding domain binding |

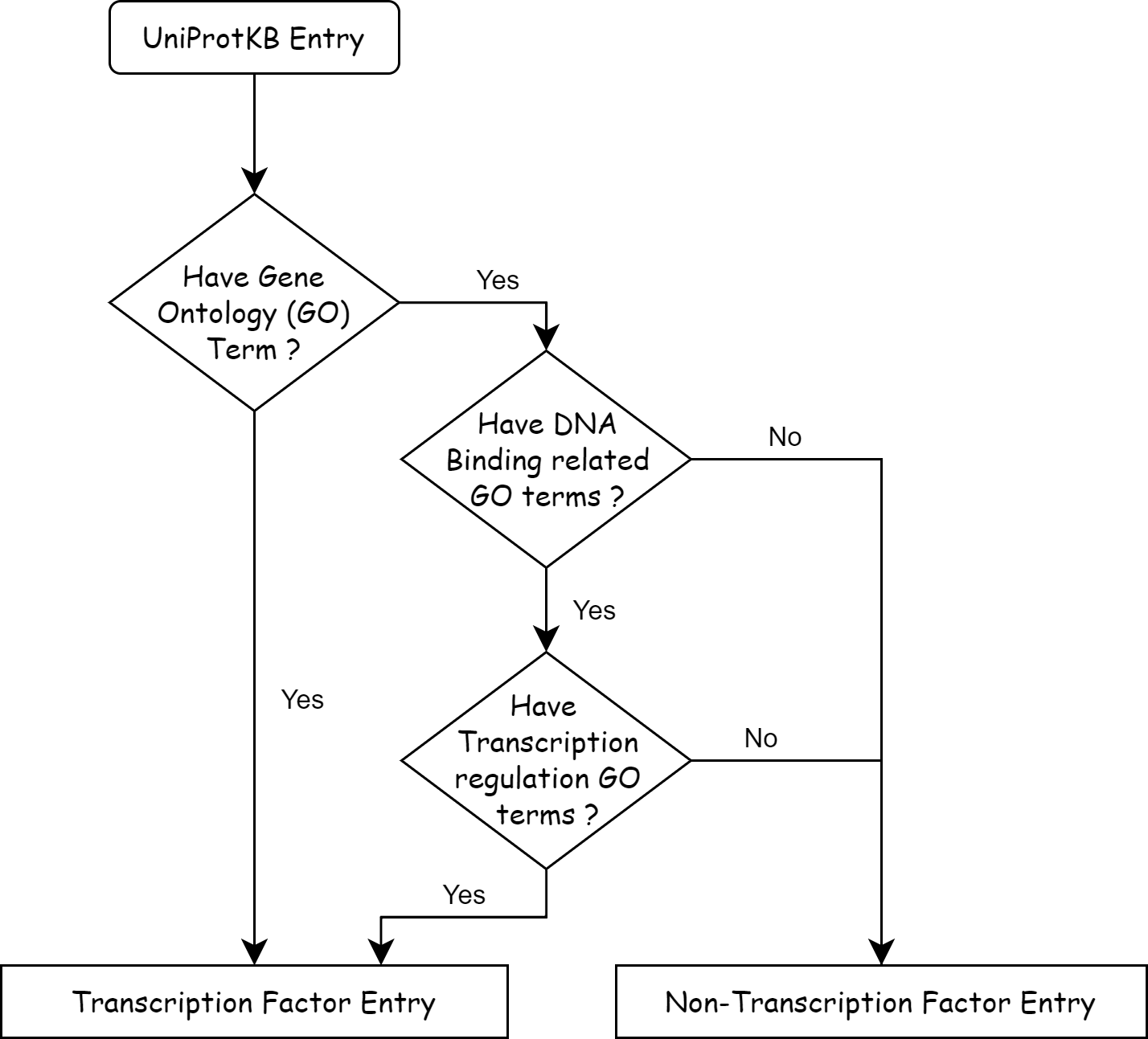
